# MRI Radiomic Signature of White Matter Hyperintensities Is Associated with Clinical Phenotypes

**DOI:** 10.1101/2021.01.24.427986

**Authors:** Martin Bretzner, Anna K. Bonkhoff, Markus D. Schirmer, Sungmin Hong, Adrian V. Dalca, Kathleen L. Donahue, Anne-Katrin Giese, Mark R. Etherton, Pamela M Rist, Marco Nardin, Razvan Marinescu, Clinton Wang, Robert W. Regenhardt, Xavier Leclerc, Renaud Lopes, Oscar R. Benavente, John W. Cole, Amanda Donatti, Christoph J. Griessenauer, Laura Heitsch, Lukas Holmegaard, Katarina Jood, Jordi Jimenez-Conde, Steven J. Kittner, Robin Lemmens, Christopher R. Levi, Patrick F. McArdle, Caitrin W. McDonough, James F. Meschia, Chia-Ling Phuah, Arndt Rolfs, Stefan Ropele, Jonathan Rosand, Jaume Roquer, Tatjana Rundek, Ralph L. Sacco, Reinhold Schmidt, Pankaj Sharma, Agnieszka Slowik, Alessandro Sousa, Tara M. Stanne, Daniel Strbian, Turgut Tatlisumak, Vincent Thijs, Achala Vagal, Johan Wasselius, Daniel Woo, Ona Wu, Ramin Zand, Bradford B. Worrall, Jane Maguire, Arne Lindgren, Christina Jern, Polina Golland, Grégory Kuchcinski, Natalia S. Rost on behalf of the MRI-GENIE and GISCOME Investigators and the International Stroke Genetics Consortium

## Abstract

**Introduction:** Neuroimaging measurements of brain structural integrity are thought to be surrogates for brain health, but precise assessments require dedicated advanced image acquisitions. By means of describing the texture of conventional images beyond what meets the naked eye, radiomic analyses hold potential for evaluating brain health. We sought to: 1) evaluate this novel approach to assess brain structural integrity by predicting white matter hyperintensities burdens (WMH) and 2) uncover associations between predictive radiomic features and patients’ clinical phenotypes.

**Methods:** Our analyses were based on a multi-site cohort of 4,163 acute ischemic strokes (AIS) patients with T2-FLAIR MR images and corresponding deep-learning-generated total brain and WMH segmentation. Radiomic features were extracted from normal-appearing brain tissue (brain mask–WMH mask). Radiomics-based prediction of personalized WMH burden was done using ElasticNet linear regression. We built a radiomic signature of WMH with the most stable selected features predictive of WMH burden and then related this signature to clinical variables (age, sex, hypertension (HTN), atrial fibrillation (AF), diabetes mellitus (DM), coronary artery disease (CAD), and history of smoking) using canonical correlation analysis.

**Results:** Radiomic features were highly predictive of WMH burden (R^2^=0.855±0.011). Seven pairs of canonical variates (CV) significantly correlated the radiomics signature of WMH and clinical traits with respective canonical correlations of 0.81, 0.65, 0.42, 0.24, 0.20, 0.15, and 0.15 (FDR-corrected p-values_CV1-6_<.001, p-value_CV7_=.012). The clinical CV1 was mainly influenced by age, CV2 by sex, CV3 by history of smoking and DM, CV4 by HTN, CV5 by AF and DM, CV6 by CAD, and CV7 by CAD and DM.

**Conclusion:** Radiomics extracted from T2-FLAIR images of AIS patients capture microstructural damage of the cerebral parenchyma and correlate with clinical phenotypes. Further research could evaluate radiomics to predict the progression of WMH.

**Research in context:** *Evidence before this study:* We did a systematic review on PubMed until December 1, 2020, for original articles and reviews in which radiomics were used to characterize stroke or cerebrovascular diseases. Radiomic analyses cover a broad ensemble of high-throughput quantification methods applicable to digitalized medical images that extract high-dimensional data by describing a given region of interest by its size, shape, histogram, and relationship between voxels. We used the search terms “radiomics” or “texture analysis”, and “stroke”, “cerebrovascular disease”, “small vessel disease”, or “white matter hyperintensities”. Our research identified 24 studies, 18 studying radiomics of stroke lesions and 6 studying cerebrovascular diseases. All the latter six studies were based on MRI (T1-FLAIR, dynamic contrast-enhanced imaging, T1 & T2-FLAIR, T2-FLAIR post-contrast, T2-FLAIR, and T2-TSE images). Four studies were describing small vessel disease, and two were predicting longitudinal progression of WMH. The average sample size was small with 96 patients included (maximum: 204). One study on 141 patients identified 7 T1-FLAIR radiomic features correlated with cardiovascular risk factors (age and hyperlipidemia) using univariate correlations. All studies were monocentric and performed on a single MRI scanner.

*Added value of this study:* To date and to the best of our knowledge, this is the largest radiomics study performed on cerebrovascular disease or any topic, and one of the very few to include a great diversity of participating sites with diverse clinical MRI scanners. This study is the first one to establish a radiomic signature of WMH and to interpret its relationship with common cardiovascular risk factors. Our findings add to the body of evidence that damage caused by small vessel disease extend beyond the visible white matter hyperintensities, but the added value resides in the detection of that subvisible damage on routinely acquired T2-FLAIR imaging. It also suggests that cardiovascular phenotypes might manifest in distinct textural patterns detectable on conventional clinical-grade T2-FLAIR images.

*Implications of all the available evidence:* Assessing brain structural integrity has implications for treatment selection, follow-up, prognosis, and recovery prediction in stroke patients but also other neurological disease populations. Measuring cerebral parenchymal structural integrity usually requires advanced imaging such as diffusion tensor imaging or functional MRI. Translation of those neuroimaging biomarkers remains uncommon in clinical practice mainly because of their time-consuming and costly acquisition. Our study provides a potential novel solution to assess brains’ structural integrity applicable to standard, routinely acquired T2-FLAIR imaging. Future research could, for instance, benchmark this radiomics approach against diffusion or functional MRI metrics in the prediction of cognitive or functional outcomes after stroke.

## Introduction

White matter hyperintensities (WMH) are a cardinal manifestation of small vessel disease (SVD).^1^ Increased WMH burden is associated with incident ischemic stroke and worse clinical outcome.^2^ Beyond ischemic stroke, WMH are also associated with vascular cognitive impairment and dementia.^3^ WMH prevalence increases with age but is also directly influenced by individual small vessel risk factors: the aggregation of cardiovascular risk factors leads to an increased WMH burden.^4^ Hence, WMH are an imaging biomarker of brain health suggestive of neurodegeneration beyond normal brain aging.^5^

Structural injury of the brain has been shown to occur at the macrostructural level, in the form of WMH, but also at the microstructural level. Advanced diffusion tensor imaging (DTI) studies have shown an age-related loss of parenchymal microstructural integrity in normal-appearing white matter (NAWM).^6^ Furthermore, perfusion-weighted imaging (PWI)-based research has also revealed age-related alterations of the blood-brain barrier with increased contrast agents’ leakage.^7^ However, such microstructural injuries are not visualized with conventional structural MRI sequences, and as DTI and PWI require special acquisition times, the outlined imaging biomarkers are not currently used in clinical routine for SVD patients. Consequently, we are in need of conventional MRI-based methodologies that better quantify SVD and brain health to ensure a widespread application and translation to clinical practice.

Radiomic analyses cover a broad ensemble of high-throughput quantification methods applicable to digitalized medical images.^8^ These methods automatically extract high-dimensional data, called radiomic features, by describing a given region of interest by its size, shape, histogram, and relationship between voxels. Because these techniques can capture slight differences in intensities and patterns that would remain undetected to a human reader, radiomics bear the potential to describe neuroimaging beyond what meets the naked eye, and thus might help to phenotype SVD.^9^ Conceivably, they may identify early underlying brain injury at the individual level with rapid clinical translatability and thus enhance personalized care in stroke and SVD.

The aim of the current study was to assess the structural integrity of the brain using a texture analysis approach and to understand the infra-radiological footprint of WMH by exploring its relationship with cardiovascular risk profiles. To do so, we analyzed 4,163 T2 FLAIR images from a large multi-site international collaborative effort studying stroke and WMH. We sought to (a) build a robust radiomic signature of the subvisible manifestations of WMH and (b) to apply canonical correlation analysis (CCA) to investigate the relation between this latent textural expression in relation to sociodemographic information and cardiovascular risk factors, providing a potentially novel approach to improve SVD and stroke care.

## Method

### Participants

We reviewed all stroke patients included in the MRI-GENetics Interface Exploration (MRI-GENIE) study, a large international multi-site collaboration of 20 sites gathering clinical, MRI imaging, and genetic data, built on top of the NINDS Stroke Genetics Network (SiGN) study. Both study design, data collection protocols, and populations have been previously described.^10^

### Ethics

The MRI-GENIE project has been approved by the MGH Institutional Review Board (IRB, Protocol #: 2001P001186 and Protocol #: 2003P000836), as well as ethics boards of the collaborating institutions. All participants or health care proxy provided signed informed consent.

### Data Collection and neuroimaging pre-processing

Clinical data were acquired within the SiGN study and comprised information on age, sex, hypertension (HTN), history of smoking, diabetes mellitus (DM), atrial fibrillation (AF). Among the 6,627 patients included across 20 sites, FLAIR images were available for 6,389 patients. Axial T2-FLAIR images were acquired between 2003 and 2011 within 48 hours of the initial stroke. They had a mean in-plane resolution of 0.7 mm (range: 0.3 mm-1.0mm) and a through-plane resolution of 6.2 mm (range: 3.0 mm-30.0 mm). Total brain, ventricle, and WMH segmentations were accomplished using deep learning methods described in detail previously.^11,12^ Briefly, total brain segmentation was done using a tailored 2D-convolutional neural network for clinical T2-FLAIR data. T2-FLAIR image intensities were normalized and scaled. Successively, WMH and ventricles were automatically segmented using distinct convolutional neural network frameworks. A total of 1,353 patients was excluded after final quality control of all T2-FLAIR images and respective segmentations; this control process is described in great detail in a previous publication.^11^ To capture the underlying processes of SVD in brain parenchyma not overtly affected by WMH, we computed masks for normal-appearing brain parenchyma by subtracting ventricles and WMH masks from total brain masks, resulting in 5,031 masks. Among those 5,031 patients, 868 were excluded for missing clinical data. As a result, a total of 4,163 patients were included across 17 different sites.

### Radiomic feature extraction

Radiomic features were extracted using the open-source toolbox PyRadiomics V2.2.0. The full list of the radiomics extraction parameters can be found online at https://github.com/MBretzner/WMH_radiomicSign

Briefly, to account for the discrepancy in voxel sizes and to reduce unwanted variance that could be originating from differences between centers and scanners, all features were extracted in-plane from down-sampled 1×1×6 mm T2-FLAIR images. Quantization was set to a fixed bin width of 5. Features extraction was performed outside of WMH on native and pre-filtered images. Filters included Laplacian of Gaussian (LoG) filters (with sigmas of 1, 2, 3 mm), wavelet decompositions, and 2D Local Binary Patterns (2D-LBD). For each patient, 118 features were computed including mask statistics, shape features, first-order histogram statistics, GLCM (Gray Level Co-occurrence Matrix) features, GLRLM (Gray Level Run Length Matrix), GLDM (Gray Level Dependence Matrix), and NGTDM (Neighboring Gray Tone Difference Matrix) features. Exhaustive and didactic descriptions and formulas of every radiomic feature and filter can be found online at https://pyradiomics.readthedocs.io/en/latest/features.html. As a result, we extracted 763 rotation invariant radiomic features per patient.

### Machine learning approach to build the radiomic signature of the WMH

To account for cerebral size differences, each WMH volume was divided by the corresponding brain volume to obtain a percentage of WMH per total brain volume. As the resultant distribution was highly skewed, it was transformed using a Box-Cox transform and is referred to as “WMH burden” in the next paragraphs.

To address the high dimensionality of the data, prediction of the WMH burden was done using an ElasticNet linear regressor. Since ElasticNet coefficient estimates are not scale-invariant, we standardized predictors, i.e., radiomics variables, to be 0 centered and have variances of the same order.

Radiomics-based predictions of WMH burden were performed in a 30-times repeated nested leave-one-out five-fold stratified cross-validation scheme, resulting in a total of 24,990 out-of-sample predictions. Predictions were plotted against ground truth values, and R2 values were computed with standard deviation.

To better understand the role of each class of radiomics and to rule out an association based solely on the size of the extraction mask, an ancillary prediction of the WMH burden was performed using only the radiomics features that only reflected the size and the shape of the analyzed brains.

The shrinkage ability of the ElasticNet regressor was leveraged to select the most predictive features of the WMH burden. The radiomic signature of the WMH was built with the features that were consistently selected across each of the 30 repetitions and therefore represented the most robust and stable predictors of WMH burden.

### Understanding the textural footprint of clinical phenotypes

Association of clinical variables and the radiomic signature of WMH burden was done via CCA, which allows studying two multivariate variable sets concomitantly.^13,14^ Indeed, traditional analyses explore relationships between many to one variable, whereas CCA can study complex many-to-many correlations, truly leveraging the power of multivariate datasets. CCA can be conceived as similar to principal component analyses in the way that each side of the data (here clinical and radiomics) undergoes a factorization into a latent representation of the variables, called canonical variates. The canonical correlation score of a canonical function represents the correlation between the two canonical variates that composes it. To extract each canonical function, CCA finds combinations of factors of the two sets so that they are maximally correlated. Canonical loadings represent the correlations between variables and their latent representation (canonical variates) and can be interpreted as the relative contribution of variables to the variates: a variable with a large loading has more impact on a variate than a variable with a smaller loading.

Radiomic features and continuous clinical variable (age) normality was assessed using the Shapiro-Wilks test and, if needed, were transformed using the R toolbox *BestNormalize*.^15^ Significance of canonical correlations was determined via permutation testing (1,000 permutations) and assessed using Wilks’ Lambda computed with Rao’s F-approximation, p-values were corrected for multiple testing with Benjamini-Hochberg procedure.^13,14^ Explained variances of the canonical functions were calculated and figured in a scree plot. Loadings were calculated to discover and characterize the impact of clinical and radiomic features on each canonical functions and thus to provide support for the interpretation of the relationship between the radiomics and clinical domains.

Overall, the goal of CCA is to find underlying representations that best describe the correlations between the two multi-dimensional datasets. Thus, this technique permits the estimation of the sources of maximal covariance between the clinical and the radiomics domains, highlighting the subvisible contribution of cardiovascular risk factors to T2 FLAIR imaging.

### Code availability

Radiomic features extraction, feature selection, and machine learning analyses were performed in python 3.7.6 using the toolbox *scikit-learn*.^16^ Canonical correlation analysis was performed in R V1.3.1056 using the toolboxes *CCA, vegan*.^17,18^ The complete codes used to perform the radiomics extraction as well as the extraction parameters and the data analysis are available here: https://github.com/MBretzner/WMH_radiomicSign

## Results

### Population

All patients included in MRI-GENIE have suffered an ischemic stroke. Population demographics are shown in **table 1**. The mean age was 62.8, and there were 42% females, median WMH volume was 4.2mL (interquartile range (IQR): 1.4-11.2). Admission NIH stroke scale (NIHSS) was available for 2,234 (53.7%) patients; median NIHSS was 3 (IQR: 1-6).

**Table 1:** Demographic and clinical characteristics of the study population (n=4,163)

|  |  |  |
| --- | --- | --- |
| **Age** | mean (SD) | 62.8 (15.0) |
| **Female** | n (%) | 1,748 (42.0 %) |
| **Hypertension** | n (%) | 2,825 (67.9%) |
| **Diabetes mellitus** | n (%) | 687 (16.5%) |
| **Atrial Fibrillation** | n (%) | 595 (14.3%) |
| **Coronary artery disease** | n (%) | 772 (18.5%) |
| **History of smoking** | n (%) | 1,331 (32.0%) |
| **Prior Stroke** | n (%) | 539 (12.9%) |
| **WMH volume** | median (IQR) | 4.2 mL (1.4 - 11.2) |
| **NIHSS\*** | Median (IQR) | 3 (1 - 6) |

Legend: NIHSS, NIH Stroke Scale; \*NIHSS was available for 2,234 (53.7%) patients included.
“Prior stroke” refers to a stroke preceding the one that led to the inclusion in the study.

### Building the latent radiomic signature of the WMH burden

The coefficient of determination of the repeated out-of-sample cross-validated predictions of the WMH burden was R2= 0.855 ± 0.011 (**figure 1)**. The average (SD) number of selected features per repetition was 150.3 (5.6). These features represented the most relevant ones in the prediction of WMH burden. To reduce the redundancy and multicollinearity of radiomic features, we built a signature of the WMH burden by only including the features that were systematically selected in every repetition. This step resulted in the automatic selection of 68 features, which are referred to as the “radiomic signature of WMH”. These features are listed in **supplementary table 1**.

**Figure 1:**
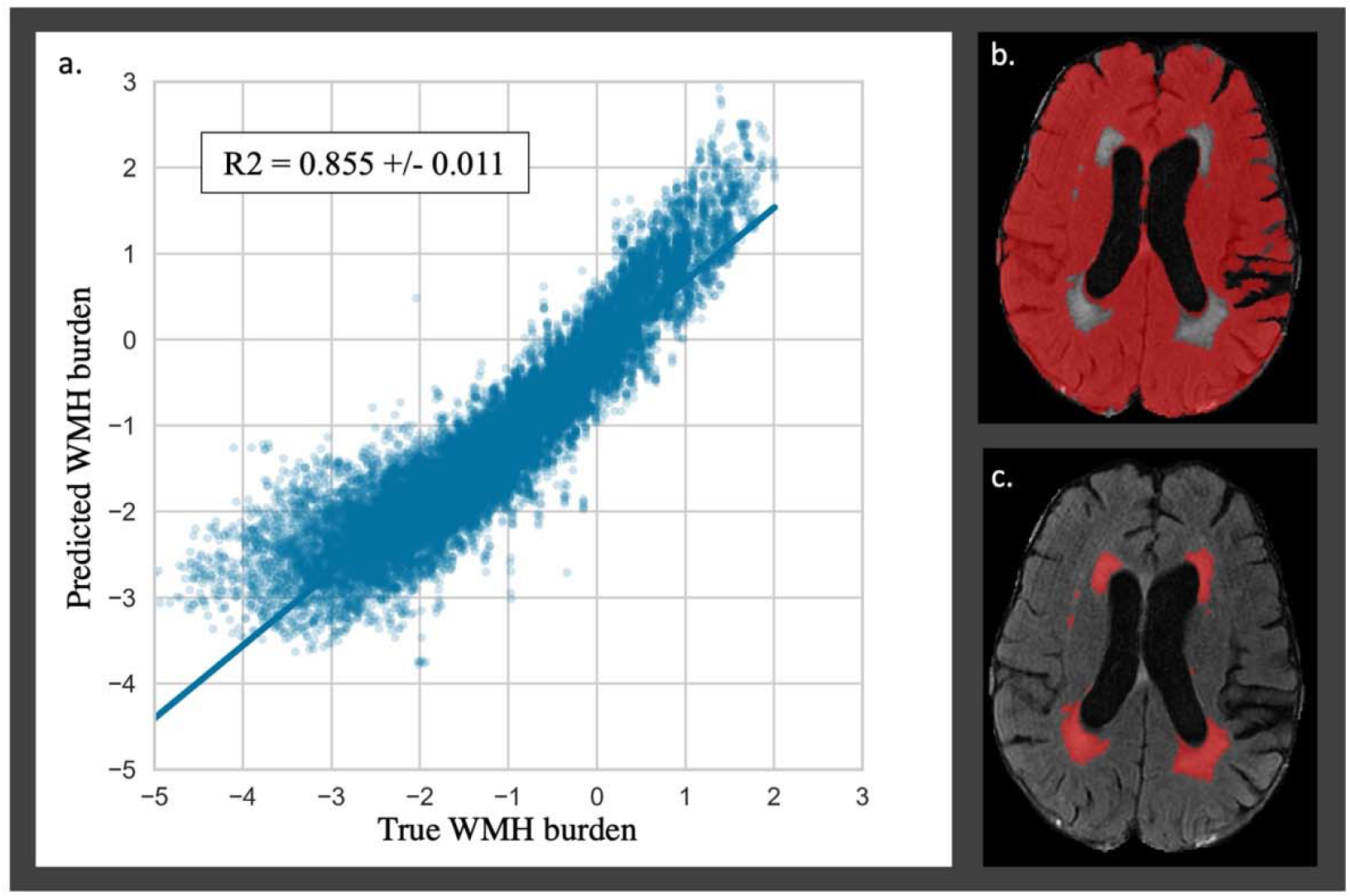
Repeated out-of-sample cross-validated predictions of WMH burden a. Predictions of the WMH burden resulted in a coefficient of determination of R2= 0.855 ± 0.011. Predicted and true WMH burdens show negative values due to the Box-Cox transformation of the WMH burden distribution. The right panels provide an illustrative example of a radiomics extraction mask (b.) and a WMH mask (c.).

Prediction performance of the WMH burden using radiomics that only describe the shape and size of the extraction mask but not voxel intensities was substantially lower with an R^2^ of 0.41 ± 0.03.

### Clinical phenotypes captured by radiomics

Aiming to discover possible links between clinical phenotypes and textural features of the radiomic signature of WMH burden, we performed a canonical correlation analysis. The CCA could identify seven canonical functions (CF 1-7) correlating the radiomics with clinical variates. All 7 canonical functions were significant (False discovery rate corrected p-values CF_1-6_ <10^−3^; CF_7_ = 0.012) with respective canonical correlations of 0.81, 0.65, 0.42, 0.24, 0.20, 0.15, and 0.15. **Figure 2** contains the scree plot of the explained variance of each CF and the correlation plot of the clinical and radiomic variates of the first canonical function with patients points colored according to their age. Loadings of the clinical and the five most impactful radiomic variables (highest loadings) of the first two canonical functions are reported in **table 2**. The bi-loading plot in **figure 3** provides a graphical interpretation relationship between the most impactful variables of the first two canonical functions. Loadings of the clinical variate of all canonical functions are shown in **table 3**, loadings of the radiomics variate are presented in **supplementary table 1**. Variables that share the same direction along a given function have a positive covariance, whereas variables that show opposing directions have negative covariance. The magnitude of the loading reflects the strength of the association.

**Figure 2:**
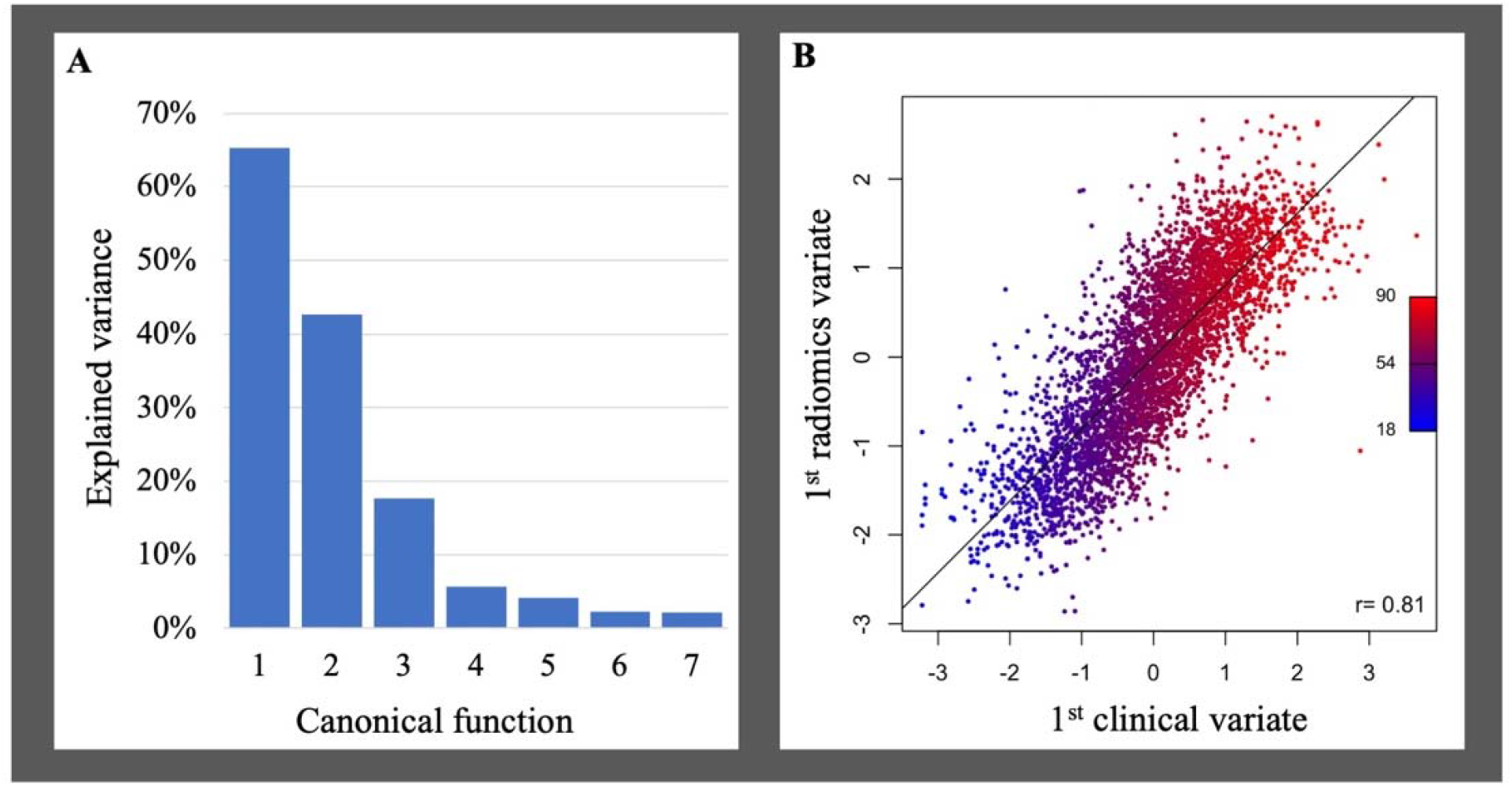
Scree plot of the explained variance per canonical function and correlation plot between the first clinical and radiomic variates. **A** Scree plot of the explained variance by canonical function. **B** Correlation plot of the first clinical and radiomics canonical variates. Each dot represents a patient and is colored according to age. The first canonical function mainly represented age. There was a very strong correlation between the clinical and the radiomics variates of *r*=0.81.

**Table 2:**
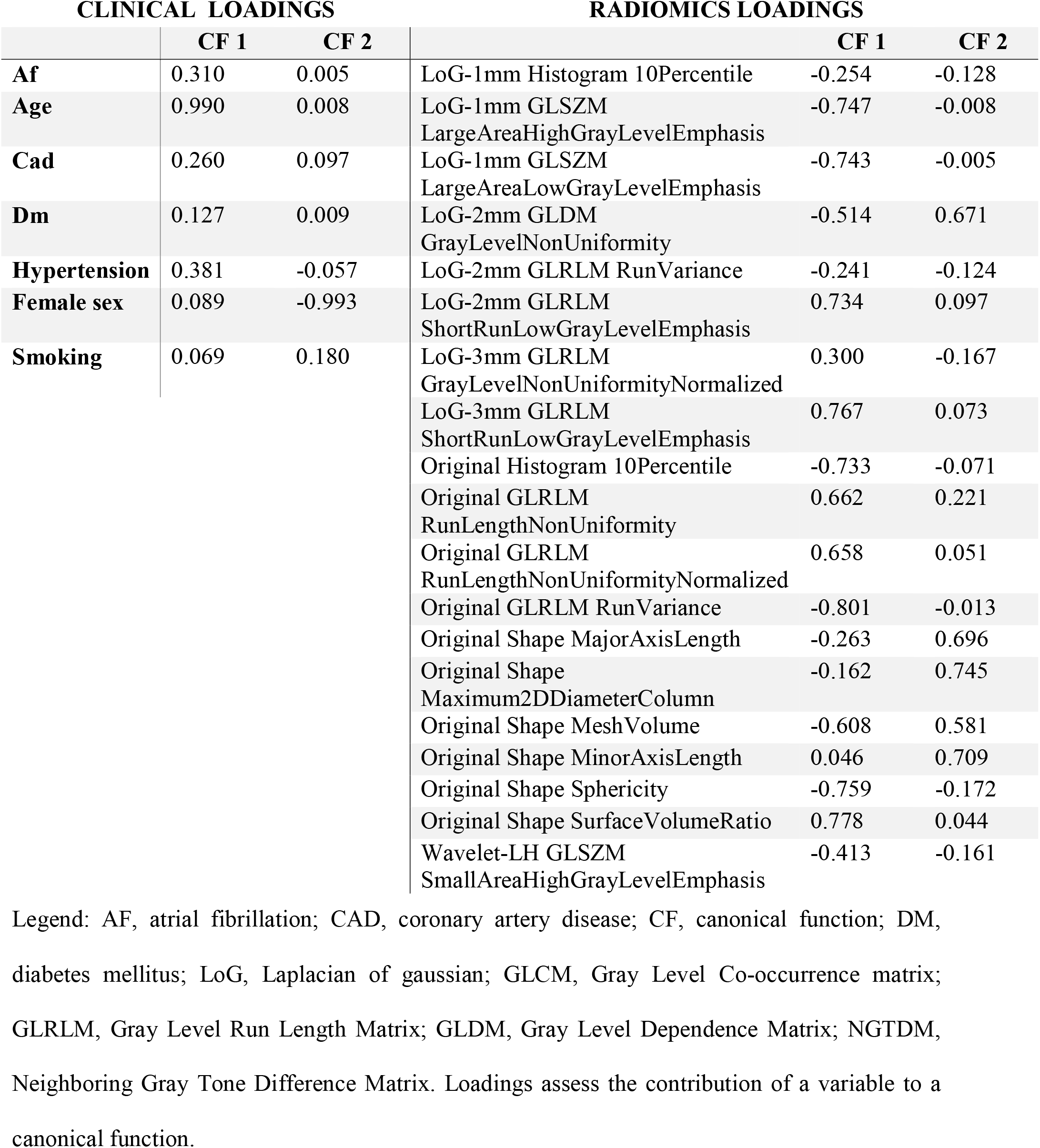
Clinical and most impactful radiomic loadings of the first two canonical functions

**Figure 3:**
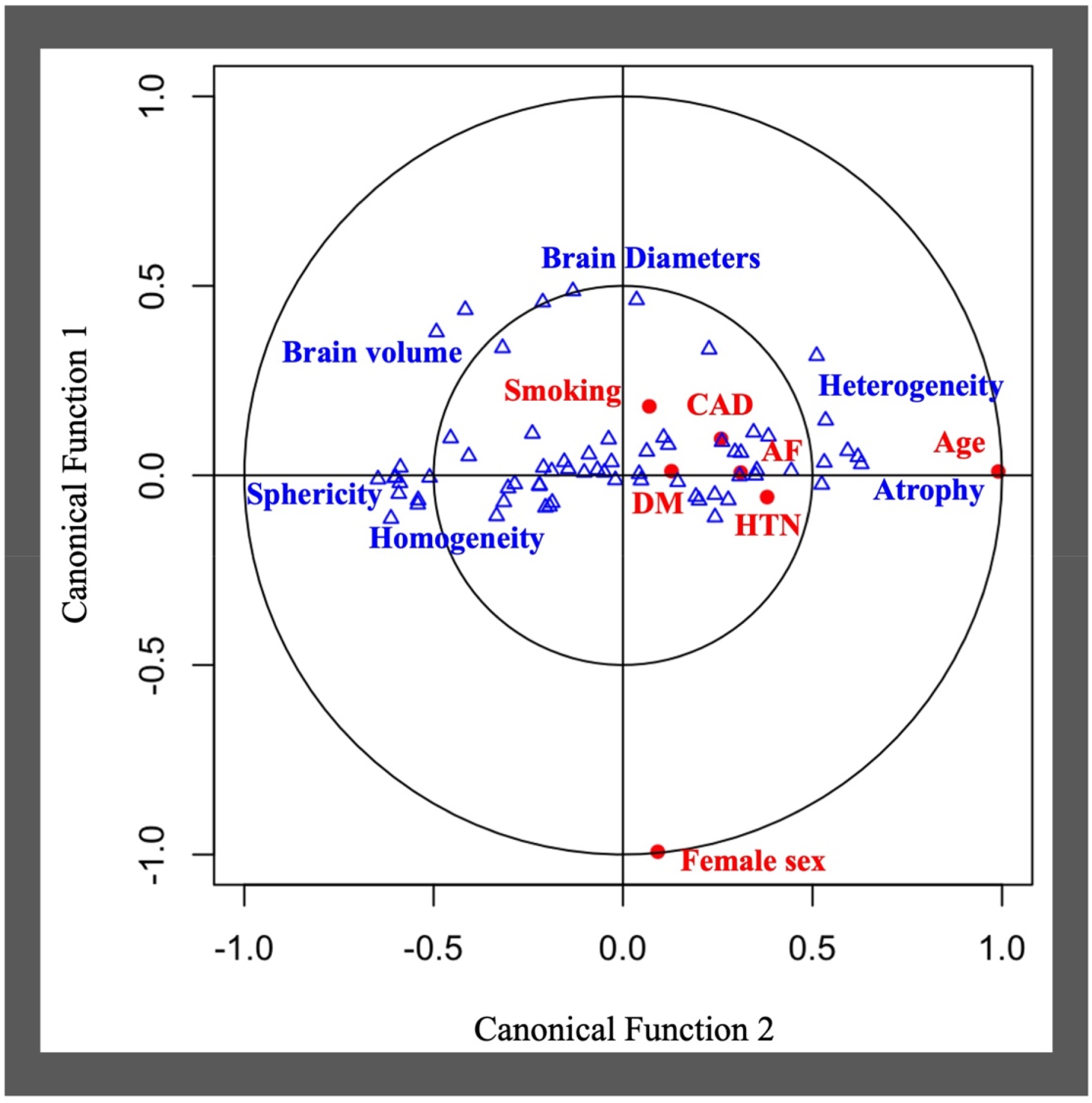
Bi-loading plot of the first two canonical variates. Clinical variables (red dot) and radiomic features (blue triangle) are positively correlated if close or negatively correlated if diagonally opposed. Blue tags were added next to correlated radiomic features representing shared textural concepts. On T2 FLAIR images, younger patients had larger brains and more homogeneous brain tissue than older patients.

**Table 3:** Clinical loadings for all seven canonical functions

| Canonical function | Clinical loadings |  |  |  |  |  |  |
| --- | --- | --- | --- | --- | --- | --- | --- |
|  | 1 | 2 | 3 | 4 | 5 | 6 | 7 |
| **Af** | 0.310 | 0.005 | 0.102 | 0.020 | 0.313 | -0.596 | 0.663 |
| **Age** | 0.990 | 0.008 | 0.096 | 0.080 | -0.045 | 0.034 | 0.017 |
| **Cad** | 0.260 | 0.097 | 0.169 | -0.157 | -0.214 | -0.744 | -0.521 |
| **Dm** | 0.127 | 0.009 | -0.443 | -0.117 | 0.781 | -0.050 | -0.401 |
| **Hypertension** | 0.381 | -0.057 | 0.067 | -0.916 | 0.055 | 0.058 | 0.030 |
| **Female sex** | 0.089 | -0.993 | -0.015 | 0.056 | -0.004 | -0.030 | 0.039 |
| **Smoking** | 0.069 | 0.180 | -0.903 | -0.055 | -0.347 | -0.132 | 0.081 |

Legend: AF, atrial fibrillation; CAD, coronary artery disease; DM, diabetes mellitus. A high loading coefficient implies a higher contribution to a canonical function. For instance, age was the most important variable when establishing the clinical variate of the first canonical function.

## Discussion

Radiomic features, extracted outside of the visible WMH, captured latent characteristics of WMH and could accurately predict WMH burden. Upon further analysis, these radiomics were associated with clinical traits relevant to WMH, such as age, sex, hypertension, history of smoking, diabetes mellitus, and coronary artery disease. Therefore, the methods presented here provide new tools to help to understand and quantify the microstructural portion of the parenchymal deterioration due to SVD in stroke and give a radiological snapshot of brain health. Importantly, our analyses relied on basic T2-FLAIR images, as commonly acquired in clinical routine and thus do not require any advanced, more costly additional imaging sequences.

WMH represent a cardinal feature among radiological manifestations of brain aging and SVD. However, DTI-^19^ and PWI-based studies suggested^20^ that WMH represent an end-stage macrostructural injury, embodying a surreptitious disease altering brain parenchyma. Our results support the hypothesis of WMH penumbra in cerebral SVD with a continuum between visible and invisible parenchymal damage.^1,21^ A major caveat of traditional advanced imaging biomarkers is their acquisition. Indeed, DTI sequences are rarely acquired routinely because of long scanning times, and PWI necessitates the injection of Gadolinium-based contrast agents. In contrast, our method can capture parenchymal microstructural integrity and hence, promises to replace additional dedicated imaging as a candidate approach to follow-up SVD progression in the clinic.

By means of our canonical correlation analysis, we estimated the associations between the radiomic signature of WMH and SVD risk factors. The influence of cardiovascular risk factors on brain tissue was previously investigated in neuropathology and advanced imaging studies yet was rarely described by analyzing the texture of conventional imaging.^1,22^ Our work complement and support previous studies on MRI textural analysis applied to SVD by *Valdes Hernandez et al*.^23^ on gadolinium-enhanced T2-FLAIR, *Bernal et al*.^24^ on dynamic spectral gadolinium-enhanced T1 weighted imaging, *Tozer and al*.^25^ on T1 and T2-FLAIR cognitive textural biomarkers, and *Shu and al*.^26^ and *Shao and al*.^27^ who could predict the progression of WMH using radiomics extracted from respectively T1-FLAIR and T2-FLAIR images. Our analyses were based on a large collection of clinical T2-FLAIR images, a routine MRI sequences acquired during both acute screening and follow-up of patients with stroke and cerebrovascular disease. Therefore, it argues for the overall clinical relevance of radiomics in stroke and SVD.

Age was the clinical aspect correlating most strongly with the radiomic signature of WMH burden and is a well-established predictor of WMH.^10,28^ Similarly, blood-brain barrier studies using PWI highlight an age-associated increased leakage of contrast agents within WMH, but also beyond, in NAWM, showcasing a possible preclinical pathogenic step leading to cognitive decline.^29^ Our findings also suggest the presence of age-related subvisible abnormalities that can notably be quantified on structural T2-FLAIR images. Radiomic features interpretation showed that decreased brain size and lower sphericity, both affected by cerebral atrophy, alongside with T2-FLAIR higher intensities and heterogeneity, were the most strongly correlated with age. On the first canonical function, age was the main variable, however, HTN, AF, and CAD were also moderately represented, painting the picture of vascular pathological brain aging. The heterogeneity and hyperintensities of the parenchyma could have maybe captured lacunes, enlarged perivascular spaces, or microbleed, which are, along with WMH, radiological hallmarks of SVD.^1^ Radiomics presented here could therefore portray a representation of a pathological brain aging process in stroke patients, depicting atrophic and heterogeneous parenchyma.

The second canonical function illustrated sex differences in tissue aspects in T2-FLAIR. The association of the radiomic signature with sex was mainly driven by shape radiomics capturing differences in brain size. This finding remains, however, independent from age-related atrophy since canonical functions analyze the unexplained variance from the previous function. Nevertheless, the female sex was also associated with greater linear edge density (GLRLM after Laplacian of Gaussian filtering), which might indicate some sex-specific textural differences in the loss of microstructural integrity, as suggested in DTI with previous findings reporting sex-specific fractional anisotropy values.^30^

The third canonical function captured a profile representing mainly patients with a history of smoking, and, to a lesser extent, diabetes, which shared common textural features describing more high spatial frequency changes in intensities which could represent diffuse and fine heterogeneity throughout the brain.

The fourth canonical function characterized a specific relation between hypertension and some textural features highlighting inhomogeneity on a lower spatial frequency after wavelet decomposition, thus describing a patchy texture. Since no other cardiovascular risk factor was represented on this dimension, it describes an age-independent specific textural manifestation of hypertension on T2-FLAIR.

Diabetes mellitus was mainly represented in the fifth dimension, correlating with textural features that illustrated overall less hyperintense parenchyma and especially those obtained after filtering with Laplacian Of Gaussian filters. Since those filters are known to act as blob detectors, they potentially captured isolated islets of damage.

The sixth canonical function related the presence of CAD and AF to a more homogeneous texture, which was, however, combined with a high impulse response to the Laplacian Of Gaussian filters of 1, 2, and 3mm sigma that could signify the presence of spots of subvisible damage of varying size of presupposed embolic origin. On the contrary, the seventh canonical function pictured the differences separating AF from CAD and DM patients, where AF patients seemed to exhibit more patches of high spatial frequency intensity changes, which could represent zones of subtly lesioned brain.

Diabetes and atrial fibrillation were represented by several dimensions meaning that the diseases in question could manifest in several distinctive aspects or stages in our data. Conditional factors that could influence such diversity in presentations include the relative control of disease by treatment or lifestyle, the patient’s stage of disease severity, genetic predispositions, and endophenotypes of varying severity.

As with any work on radiomics, the main pitfall remains the *curse of dimensionality*, which refers to a very high number of variables. Consequently, one of the strengths of our study was the available sample size, allowing us to truly leverage both machine learning methodologies and multivariate modeling to select and characterize relevant radiomic variables in a data-driven fashion. In fact, to date and to the best of our knowledge, this is the largest radiomics study performed on any topic. Previous work on radiomics of SVD studied smaller datasets (<250 participants) and thus did not permit powerful unsupervised feature selections.^23–25^ Another added value of the present study is its multicentric design. Our study is the first to explore radiomics of SVD in a large and multicentric cohort. By implementing multiple measures, such as down-sampling and intensity normalization, to prevent differences originating from acquisition parameters discrepancies, we could reach homogeneous results across all centers while capturing relevant sources of variance, as depicted by the low error of our WMH burden predictions. Another source of unwanted variance in radiomics analyses is segmentation, however, we here built upon previous results obtained with state-of-the-art deep learning-based, fully automated segmentation methods that could produce consistent outlines of brains, WMH, and ventricles from T2-FLAIR.^11,12^ Preventive measures we implemented, especially down-sampling and intensity normalization, may have come at the cost of losing pertinent information. However, that impact might have been mitigated thanks to our large sample size. We thus emphasize the capital importance of international collaborations, such as the MRI-GENIE consortium, to gather large datasets, especially in the era of quantitative imaging and personalized medicine.

### Limitations and future directions

We acknowledge several limitations; first, stroke lesion outlines were not available and thus not accounted for. Therefore, they could have been measured by radiomics and biased WMH predictions, e.g., caused the overestimation in predictions we observed in both the lower and upper tails of the predicted WMH burden distribution. Overall, the median size of ischemic stroke lesions in this cohort is expected to be small, as the median NIHSS was 3. Moreover, the radiomic analysis conducted here provides a single value per radiomic variable per patient, averaging the textural presentation over the whole extraction zone and thus largely decreasing the impact of small lesions. Regarding large lesions, the corresponding perturbated radiomic value could have been assimilated to an outlier and then mitigated by the ElasticNet model, which includes an L1 regularization that improves its robustness to extreme values. Other SVD imaging features were also not accounted for, such as microbleeds or enlarged perivascular space, which have been previously linked to radiomic features.^23,25^

Secondly, radiomics were extracted outside of the WMH but not specifically within the white matter. Future research could evaluate the impact of co-registration and resampling on radiomics of SVD, then benchmark radiomics of NAWM against more traditional DTI metrics in the prediction of clinical outcomes and therefore provide a more straightforward method to quantify microstructural integrity.

## Conclusion

In a large cohort of ischemic stroke patients, we demonstrated that radiomic features predicted WMH burden and were associated with clinical factors. By applying machine learning methods to radiomics analyses of T2-FLAIR images from a large multi-site ischemic stroke cohort, we could characterize the latent expression of small vessel disease that extends beyond the visible WMH and subsequently uncover links associating cardiovascular risk factors to distinct textural patterns. Radiomics analysis may hold promise to become a cost-effective tool to quantify microstructural damage on routinely acquired images in the follow-up of SVD and stroke patients, once externally validated.

### Data sharing

Upon reasonable request to the corresponding author and pending approval from local IRBs, data will be made available to replicate results presented in this manuscript.

## Acknowledgment statement

(including conflict of interest and funding sources):

The authors have declared that no competing interests exist.

The MRI-GENIE study was funded by NIH NINDS (R01NS086905).

MB was supported by the ISITE-ULNE Fundation, the Société Française de Neuroradiologie, the Société Française de Radiologie, the Thérèse and René Planiol Fundation. MRE is supported by the American Academy of Neurology and MGH Executive Council on Research. TT was supported by the Helsinki University Central Hospital, Sigrid Juselius Foundation, Sahlgrenska University Hospital, and University of Gothenburg. PMR is supported by NIH K01 HL128791. The funders had no role in study design, data collection and analysis, decision to publish, or preparation of the manuscript.

## Author contributions

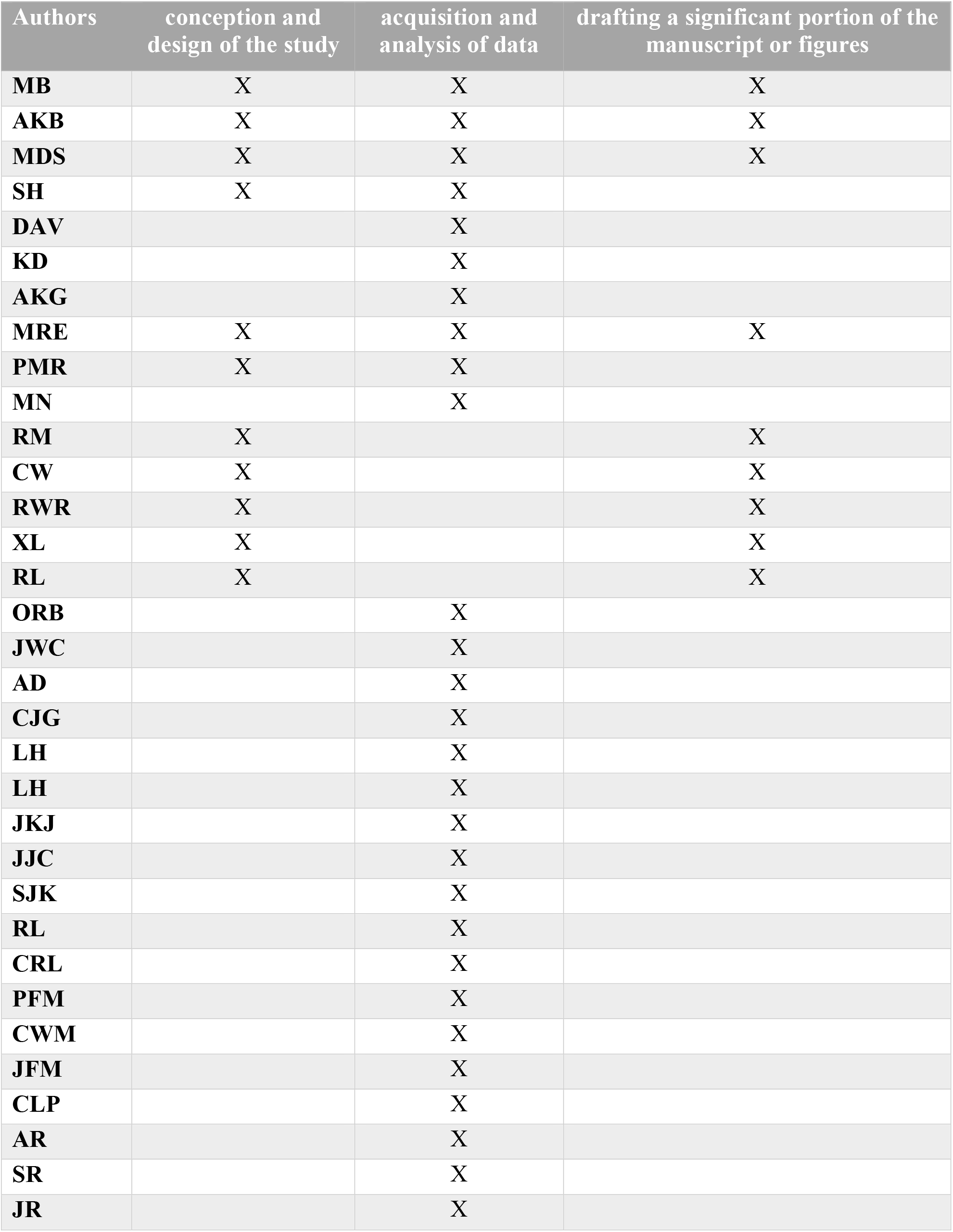
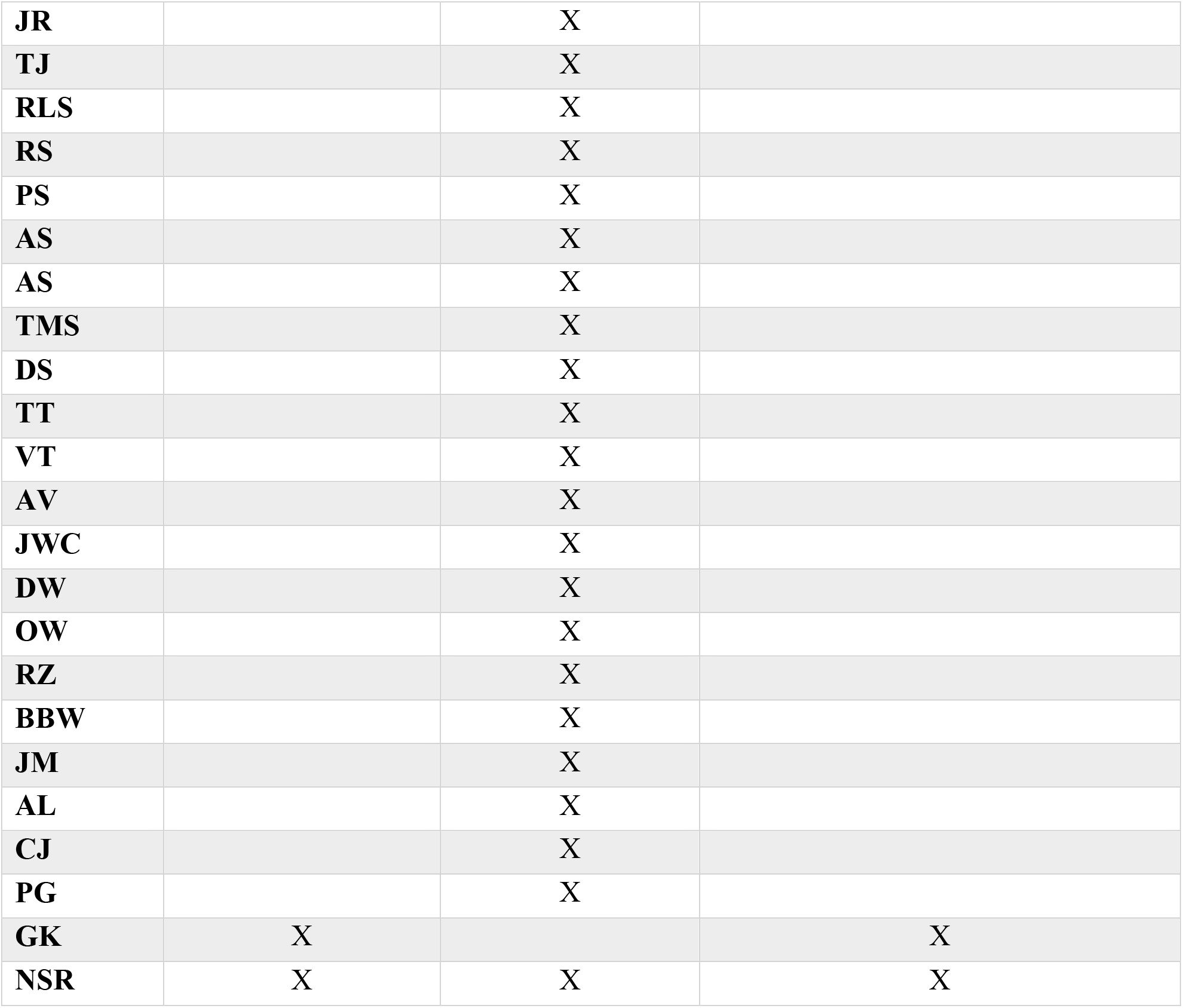

