## supplementary table 1. for "MRI Radiomic Signature of White Matter Hyperintensities Is Associated with Clinical Phenotypes"

**Supplementary table 1: Radiomic signature loadings.**

| Canonical function | 1 | 2 | 3 | 4 | 5 | 6 | 7 |
| --- | --- | --- | --- | --- | --- | --- | --- |
| Mask original Voxel Number | -0·111 | 0·087 | 0·416 | 0·134 | 0·056 | -0·081 | -0·171 |
| LBP Histogram Mean | -0·233 | 0·018 | 0·134 | -0·191 | -0·301 | 0·020 | -0·026 |
| LBP GLCM Correlation | 0·053 | 0·006 | -0·277 | 0·057 | 0·225 | 0·013 | 0·084 |
| LBP GLCM Imc1 | 0·005 | -0·028 | -0·110 | 0·095 | -0·203 | 0·210 | 0·134 |
| LBP GLRLM GrayLevelVariance | -0·084 | 0·024 | 0·292 | -0·137 | -0·274 | 0·005 | -0·099 |
| LoG-1mm-Histogram 10Percentile | -0·254 | -0·128 | -0·176 | 0·013 | 0·192 | 0·162 | 0·067 |
| LoG-1mm-Histogram 90Percentile | 0·133 | 0·153 | -0·144 | -0·094 | -0·041 | -0·282 | 0·073 |
| LoG-1mm-Histogram Median | -0·632 | -0·007 | -0·421 | -0·103 | 0·011 | 0·134 | 0·175 |
| LoG-1mm-Histogram Skewness | 0·251 | -0·100 | 0·155 | -0·082 | 0·005 | -0·046 | -0·219 |
| LoG-1mm-GLDM DependenceEntropy | -0·047 | 0·146 | -0·074 | -0·093 | 0·123 | -0·235 | 0·088 |
| LoG-1mm-GLRLM HighGrayLevelRunEmphasis | -0·504 | 0·079 | -0·444 | -0·181 | 0·020 | 0·048 | 0·218 |
| LoG-1mm-GLSZM LargeAreaHighGrayLevelEmphasis | -0·747 | -0·008 | -0·236 | 0·000 | -0·063 | -0·026 | 0·178 |
| LoG-1mm-GLSZM LargeAreaLowGrayLevelEmphasis | -0·743 | -0·005 | -0·127 | 0·062 | -0·056 | 0·041 | 0·139 |
| LoG-1mm-GLSZM SmallAreaHighGrayLevelEmphasis | 0·300 | -0·078 | 0·567 | 0·088 | -0·094 | -0·027 | -0·176 |
| LoG-1mm-GLSZM SmallAreaLowGrayLevelEmphasis | -0·273 | -0·041 | 0·055 | -0·029 | -0·182 | 0·024 | 0·115 |
| LoG-2mm-Histogram 10Percentile | -0·354 | -0·034 | -0·329 | -0·070 | 0·191 | 0·157 | 0·021 |
| LoG-2mm-Histogram RootMeanSquared | 0·385 | 0·091 | 0·223 | 0·109 | -0·141 | -0·162 | -0·006 |
| LoG-2mm-GLCM ClusterShade | 0·649 | -0·038 | 0·334 | 0·130 | -0·031 | 0·060 | -0·243 |
| LoG-2mm-GLDM GrayLevelNonUniformity | -0·514 | 0·671 | -0·096 | 0·013 | -0·064 | 0·043 | 0·052 |
| LoG-2mm-GLRLM HighGrayLevelRunEmphasis | -0·563 | 0·153 | -0·007 | -0·059 | 0·328 | 0·168 | 0·012 |
| LoG-2mm-GLRLM RunLengthNonUniformityNormalized | 0·427 | 0·173 | 0·193 | -0·088 | -0·034 | 0·028 | -0·083 |
| LoG-2mm-GLRLM RunVariance | -0·241 | -0·124 | -0·029 | 0·128 | -0·055 | 0·058 | -0·180 |
| LoG-2mm-GLRLM ShortRunLowGrayLevelEmphasis | 0·734 | 0·097 | 0·152 | -0·047 | -0·011 | -0·044 | -0·203 |
| LoG-2mm-GLSZM GrayLevelNonUniformity | 0·282 | 0·510 | 0·446 | 0·087 | -0·056 | -0·001 | -0·213 |
| LoG-2mm-GLSZM SmallAreaLowGrayLevelEmphasis | 0·343 | -0·100 | 0·136 | 0·013 | -0·069 | 0·004 | -0·087 |
| LoG-3mm-Histogram 10Percentile | -0·261 | 0·033 | -0·331 | -0·133 | 0·225 | 0·206 | 0·032 |
| LoG-3mm-Histogram 90Percentile | 0·366 | 0·093 | -0·049 | 0·153 | 0·145 | -0·304 | 0·107 |
| LoG-3mm-Histogram InterquartileRange | 0·386 | 0·001 | 0·197 | 0·075 | -0·053 | -0·281 | -0·043 |
| LoG-3mm-Histogram Skewness | 0·550 | 0·016 | 0·101 | 0·067 | 0·141 | -0·010 | -0·223 |
| LoG-3mm-GLDM LargeDependenceEmphasis | -0·671 | -0·114 | -0·259 | -0·018 | -0·096 | -0·039 | 0·084 |
| LoG-3mm-GLRLM GrayLevelNonUniformityNormalized | 0·300 | -0·167 | -0·179 | -0·029 | -0·374 | -0·047 | 0·168 |
| LoG-3mm-GLRLM HighGrayLevelRunEmphasis | -0·296 | 0·170 | 0·182 | 0·031 | 0·373 | 0·046 | -0·172 |
| LoG-3mm-GLRLM RunEntropy | -0·670 | -0·098 | -0·340 | -0·053 | -0·064 | -0·059 | 0·124 |
| LoG-3mm-GLRLM ShortRunLowGrayLevelEmphasis | 0·767 | 0·073 | 0·235 | 0·008 | 0·013 | 0·003 | -0·183 |
| original Histogram 10Percentile | -0·733 | -0·071 | 0·061 | -0·030 | -0·309 | 0·014 | 0·106 |
| original GLRLM RunEntropy | -0·730 | -0·028 | -0·311 | -0·084 | -0·054 | 0·004 | 0·179 |
| original GLRLM RunLengthNonUniformity | 0·662 | 0·221 | 0·324 | 0·131 | 0·069 | -0·025 | -0·194 |
| original GLRLM RunLengthNonUniformityNormalized | 0·658 | 0·051 | 0·350 | 0·166 | 0·092 | -0·061 | -0·191 |
| original GLRLM RunVariance | -0·801 | -0·013 | -0·187 | 0·077 | -0·002 | -0·039 | 0·191 |
| original GLRLM ShortRunEmphasis | 0·437 | 0·020 | 0·361 | 0·155 | 0·095 | -0·094 | -0·223 |
| original GLSZM GrayLevelNonUniformityNormalized | 0·179 | -0·026 | 0·252 | -0·087 | -0·261 | -0·173 | 0·151 |
| original GLSZM GrayLevelVariance | -0·179 | 0·026 | -0·251 | 0·090 | 0·261 | 0·174 | -0·151 |
| original GLSZM SmallAreaLowGrayLevelEmphasis | -0·025 | -0·018 | 0·005 | -0·039 | 0·126 | 0·095 | -0·306 |
| original GLSZM ZoneEntropy | 0·148 | 0·124 | 0·049 | 0·093 | -0·061 | -0·110 | 0·147 |
| original NGTDM Complexity | -0·038 | 0·054 | -0·254 | 0·138 | 0·265 | 0·114 | -0·166 |
| original Shape LeastAxisLength | -0·393 | 0·517 | -0·127 | -0·146 | 0·050 | 0·062 | -0·017 |
| original Shape MajorAxisLength | -0·263 | 0·696 | -0·074 | 0·101 | -0·203 | -0·030 | -0·013 |
| original Shape Maximum2DDiameterColumn | -0·162 | 0·745 | -0·018 | -0·076 | 0·033 | -0·051 | 0·139 |
| original Shape MeshVolume | -0·608 | 0·581 | -0·175 | -0·019 | -0·031 | 0·025 | 0·105 |
| original Shape MinorAxisLength | 0·046 | 0·709 | -0·039 | 0·074 | 0·010 | 0·032 | 0·050 |
| original Shape Sphericity | -0·759 | -0·172 | -0·266 | -0·054 | -0·033 | -0·031 | 0·189 |
| original Shape SurfaceArea | 0·634 | 0·482 | 0·251 | 0·066 | 0·009 | 0·027 | -0·188 |
| original Shape SurfaceVolumeRatio | 0·778 | 0·044 | 0·262 | 0·048 | 0·034 | 0·025 | -0·193 |
| Wavelet-HH GLRLM GrayLevelNonUniformityNormalized | 0·433 | -0·001 | 0·406 | 0·197 | 0·121 | -0·038 | -0·267 |
| Wavelet-HH GLRLM RunEntropy | -0·390 | -0·105 | 0·064 | 0·074 | -0·036 | 0·044 | 0·182 |
| Wavelet-HH GLSZM LargeAreaHighGrayLevelEmphasis | -0·728 | 0·034 | -0·215 | -0·078 | -0·033 | 0·056 | 0·204 |
| Wavelet-HL Histogram Mean | -0·376 | -0·051 | -0·357 | -0·093 | -0·005 | 0·092 | 0·142 |
| Wavelet-HL Histogram Skewness | -0·232 | -0·109 | 0·172 | 0·104 | 0·003 | 0·138 | -0·114 |
| Wavelet-HL Histogram Variance | 0·078 | 0·097 | 0·032 | -0·022 | -0·281 | -0·122 | -0·032 |
| Wavelet-HL GLCM JointEntropy | -0·192 | 0·055 | -0·125 | -0·025 | 0·180 | 0·268 | 0·072 |
| Wavelet-HL GLRLM HighGrayLevelRunEmphasis | -0·127 | 0·012 | -0·057 | -0·083 | -0·252 | -0·040 | 0·115 |
| Wavelet-HL GLRLM RunVariance | -0·274 | -0·038 | -0·377 | -0·073 | 0·143 | -0·094 | 0·076 |
| Wavelet-HL GLSZM ZoneEntropy | 0·323 | 0·135 | -0·402 | -0·042 | 0·126 | -0·126 | -0·087 |
| Wavelet-LH GLCM Imc2 | -0·061 | 0·013 | 0·251 | -0·243 | -0·024 | -0·153 | -0·111 |
| Wavelet-LH GLCM MaximumProbability | 0·061 | -0·019 | 0·401 | -0·166 | 0·005 | -0·141 | -0·162 |
| Wavelet-LH GLDM DependenceVariance | 0·238 | -0·086 | 0·131 | 0·278 | 0·181 | 0·035 | 0·093 |
| Wavelet-LH GLSZM SmallAreaHighGrayLevelEmphasis | -0·413 | -0·161 | 0·318 | -0·114 | -0·160 | 0·135 | 0·124 |
| Wavelet-LH GLSZM ZoneEntropy | 0·475 | 0·156 | -0·147 | 0·157 | 0·171 | 0·006 | -0·147 |

LoG, Laplacian of gaussian; LBP, Local Binary Patter; GLCM, Gray Level Co-occurrence matrix; GLRLM, Gray Level Run Length Matrix; GLDM, Gray Level Dependence Matrix; NGTDM, Neighboring Gray Tone Difference Matrix. Loadings assess the relative contribution of a variable to a canonical function.
